# Ipflufenoquin, Novel Dihydroorotate Dehydrogenase Inhibitor: Efficacy Against Strawberry Diseases and Docking Simulations via AlphaFold2 and Autodock Vina

**DOI:** 10.1101/2024.12.17.628804

**Authors:** Sungyu Choi, Doeun Son, Hyeong-rok Jang, Soyoon Park, Haifeng Liu, Hyeon-gyeong Lee, Hyunkyu Sang

## Abstract

In 2023, fungal diseases, including *Botrytis cinerea* (gray mold), were isolated from various strawberry fields in Korea. Ipflufenoquin, a novel class 2 dihydroorotate dehydrogenase (DHODH) inhibitor, was applied to plant pathogenic fungal species. EC_50_ values indicated that ipflufenoquin achieved high sensitivity in *B. cinerea* while maintaining broad-spectrum activity against other fungi. Fruit and greenhouse trials confirmed its robust efficacy in gray mold diseases, including those resistant to conventional fungicides like pyraclostrobin, fluxapyroxad, and benomyl. However, the ipflufenoquin application was less effective against fungal pathogens like blossom blight and Rhizopus rot, causing shifts in the fungal community and allowing less sensitive species to emerge. Structural prediction using Alphafold2 and molecular docking simulation showed ipflufenoquin was bound to the quinone binding tunnel in DHODH and highlighted the correlation between protein-ligand binding affinity and fungicide susceptibility, with species showing higher affinities being more sensitive. This study is the first to report sensitivity of strawberry fungal species to ipflufenoquin, and this fungicide is recommended in strawberry farms against gray mold disease. Additionally, we conducted in silico predictions of fungicide sensitivity across various plant pathogenic fungi using Alphafod2 and Autodock Vina. This establishes a foundation for future computational analyses to predict the effectiveness of fungicides, offering important baseline data to enhance fungicide management strategies.

## 1. Introduction

Strawberry diseases significantly impact yield and quality, necessitating effective detection and management strategies. Recent research has highlighted various diseases affecting strawberries, including anthracnose, root rot, and gray mold, and explores advanced methodologies for their identification. Major strawberry diseases worldwide are: (1) ‘anthracnose’, caused by *Colletotrichum* spp., resulting in up to 70% yield loss in North America and manifesting as lesions on leaves and fruit; (2) ‘crown rot’, caused by *Fusarium* spp., which causes wilting and necrosis, threatening strawberry production; and (3) ‘gray mold’, caused by *Botrytis cinerea* which is prevalent in strawberry plantations and one of the most harmful diseases (Mykhailenko et al., 2022; Zhang et al., 2022; Rahman et al., 2023).

The necrotrophic fungus, *B. cinerea*, affects diverse plant species, including fruits, vegetables, and ornamental plants, leading to severe economic losses (Williamson et al., 2007). Understanding the biology, epidemiology, and management of *B. cinerea* is crucial for devising effective strategies to mitigate its impact on crop yields. The economic ramifications of gray mold disease, caused by *B. cinerea*, are substantial within the agricultural sector. This fungal pathogen damages crops and undermines post-harvest storage and transportation, incurring significant financial losses for growers, distributors, and consumers alike (Nakielska et al., 2024). These economic effects of *B. cinerea* underscore the importance of developing sustainable disease management practices.

Chemical control is one of the primary methods utilized to manage disease caused by *B. cinerea* in agricultural settings. Various fungicides with diverse modes of action are employed to combat this pathogen, aiming to disrupt crucial biological processes essential for its growth and reproduction. Understanding the mechanisms of action of these fungicides is pivotal for optimizing their efficacy and minimizing adverse environmental impacts (Leroux et al., 2002). Until recently, multiple resistance (MLR) and multidrug resistance (MDR) were reported by mechanism of action: succinate dehydrogenase inhibitors (SDHIs; FRAC 7), quinone outside inhibitors (QoIs; FRAC 11), benzimidazoles (MBC; FRAC 1), dicarboximides (DCs; FRAC 2), demethylation inhibitors (SBI: Class I; FRAC 3), and phenylpyrroles (PP; FRAC 12) fungicides. These resistances have significantly reduced the efficacy of these fungicides, leading to cross-resistance issues and limiting the use of fungicides with conventional modes of action (Myresiotis et al., 2007; Banno et al., 2009; Sofianos et al., 2023; Kim et al., 2023).

As a new avenue for managing fungal pathogens, the development and utilization of dihydroorotate dehydrogenase inhibitors (DHODHIs) has shown promise in combating plant diseases caused by pathogenic fungi (Higashimura et al., 2023). Various structures of quinoline compounds have shown wide biological activities, such as antibacterial, antifungal, antiviral, antimalarial, and anticancer, targeting DHODH and providing potential discovery for new pesticides (Huang et al., 2015; Du Pré et al., 2018; Boschi et al., 2019; Senerovic et al., 2019; Liu et al., 2020). These compounds bind specifically to the ubiquinone-binding tunnel of class 2 dihydroorotate dehydrogenase, preventing ubiquinone from participating in the electron transfer process necessary for the enzyme’s catalytic function. By disrupting this process, these inhibitors interfere with critical metabolic pathways, including the production of pyrimidines, necessary for fungal growth and RNA synthesis. This blockage hampers the survival and proliferation of pathogenic fungi. (Boschi et al., 2019). For example, quinofumelin, a quinoline fungicide, is used on fruit trees, leafy vegetables, rapeseed, and rice, and exhibits strong antifungal activity against mycelial growth (Higashimura et al., 2023). F901318(olorofim), a chemical DHODHI, showed potent antifungal activities on *Aspergillus* spp. and displayed application range in fungal species (Oliver et al., 2016). Chosen for this study, ipflufenoquin, with a chemical structure including a quinoline ring substituted with multiple fluorine atoms at specific positions (2, 3, 7, and 8) and a hydroxypropan-2-yl phenoxy group, was developed by Nippon Soda Co., Ltd. (Japan) and exhibits potent antifungal activity against wide plant filamentous diseases, including *Aspergillus fumigatus* (Van Rhijn et al., 2023).

Molecular docking plays a critical role in computational biology, predicting how small molecules like chemical ligands interact with target proteins (Agu et al., 2023). This method is widely used in drug discovery and has extended its applications to the agricultural sector, particularly in pesticide design (Hou et al., 2023). Docking simulations help predict how chemicals bind to proteins in pests, offering a pathway to create more effective and targeted pesticides. Recent advancements in docking algorithms have significantly improved the accuracy of predicting interactions. For example, molecular docking has been used to identify fungicides that target critical enzymes in fungal pathogens, such as *Fusarium* species, which are responsible for crop diseases (Aamir et al., 2018). Moreover, AI-powered tools, particularly those for predicting protein structures, have significantly broadened the scope of molecular docking studies (Holcomb et al., 2023). Tools like AlphaFold2 enable accurate predictions of protein structures across various species, allowing researchers to explore molecular interactions and subtle structural changes at an *in silico* level. These innovations are useful in assessing the effectiveness of fungicides by simulating their binding with the target proteins (Cen et al., 2023). This enables more detailed assessments of fungicide-protein interactions in various fungal pathogens, ultimately contributing to the development of more targeted and effective solutions for controlling plant diseases.

In this study, the application and evaluation of ipflufenoquin to strawberry fungal diseases, including *B. cinerea* strains from strawberry farms in Korea, was performed and a control efficacy test of ipflufenoquin against gray mold disease was investigated in a strawberry greenhouse. The control spectrum of ipflufenoquin was investigated using molecular docking prediction via Alphafold2 and Autodock Vina, which allowed for a detailed comparison between the computationally predicted binding affinities and the actual antifungal efficacy observed in in vitro assays.

## 2. Materials and Methods

### 2.1 Fungal isolation and identification

In 2023, 88 strains of *B. cinerea* were collected from symptomatic strawberries produced in 88 different strawberry farms at various locations in Korea (Table. S1). Each strain was obtained from single-spore isolation and was stored at -80°C in 25% glycerol solution. The identification markers of ITS region and NEP2 gene using the primer pairs ITS5/ITS4 and NEP2forD/NEP2revD were amplified by polymerase chain reaction (PCR) (Staats et al., 2007). To investigate fungal species in strawberry plants, sixty-four field isolates were collected from strawberry plants, including fruits, flowers, leaves, rhizosphere, and soil samples, from December 2022 to April 2023 in the greenhouse located in Gwangju, Korea. Each plant tissue was rinsed in sodium hypochlorite (NaClO) 1% for 1 minute and washed with distilled water three times. Surface-sterilized tissues were placed onto PDA agar plates with antibiotics and incubated at 25 °C for three days in the dark, and a single colony from each plate was isolated. Soil samples were diluted 10^3^ times with distilled water and spread onto PDA agar plates containing antibiotics (100 mg L^-1^ of ampicillin, 50 mg L^-1^ of streptomycin, and 100mg L^-1^ of spectinomycin) for fungal isolation. Fungal identification was performed morphologically, and the ITS region was amplified using primer pairs ITS5/ITS4 initially. For classification of the fungal species, amplification was performed of each region of ALT1 and EF1 for *Alternaria* spp., EF1 for *Chaetomium* spp., DHODH *for Cladosporium* spp., GAPDH for *Colletotrichum* spp., TUB2 for *Fusarium* spp., EF1 for *Trichoderma* spp. (Ahn et al., 2022; Burkhardt et al., 2019; Cai and Druzhinina, 2021; Chung et al., 2020; Dong et al., 2023; Hong et al., 2005, p. 1; Kwon et al., 2014; Lee et al., 2023; Wang et al., 2016). Additional fungal species, including *Aspergillus flavus*, *Colletotrichum truncatum*, *Pythopthora sojae*, *Rhizoctonia solani* AG1, and AG2-2-ⅢB, *Fusarium asiatichum,* were used to investigate sensitivity to ipflufenoquin. These strains had been stored in the Chonnam National University Molecular Microbiology Laboratory (CNUMML).

### 2.2 In vitro sensitivity tests of fungal strains to ipflufenoquin

In *B. cinerea*, the sensitivity to ipflufenoquin was determined by the inhibition rate of mycelium growth and conidia germination in fungicide-amended media. The strains were cultured on PDA at 25 °C for 7 days. The assay was conducted using PDA media amended with ipflufenoquin at concentrations of 0, 0.01, 0.1, 1, 10, and 100 mg L^-1^. For mycelial inoculation, the 88 strains were cultured on PDA media at 25 °C in the dark for 7 days. Five mm-agar plugs were prepared and inoculated at the center of ipflufenoquin-amended media. Conidia for germination assays were harvested from PDA media, which were exposed to near-ultraviolet light in light-dark conditions for 12h/12h, and 1×10^5^ conidia suspensions were inoculated to fungicide media. After incubation for 3 days at 25°C, the diameters of the colonies were measured using a digital caliper (Mitutoyo, Japan). The EC_50_ (effective concentration for 50% inhibition) values were calculated for each isolate using R software (v4.2.3; R Development Core Team 2023). Two replicate plates were used for each treatment, and all experiments were performed two times.

In vitro sensitivity to ipflufenoquin of other fungal pathogens, which were isolated from the greenhouse located in Gwangju, was conducted. Mycelial plugs or 1×10^5^ conidia suspension was transferred to PDA amended with ipflufenoquin at 0, 0.001, 0.01, 0.1, 1, 10, and 100 mg L^-1^. All fungal strains were incubated in the dark at 25°C for up to 9 days to ensure sufficient growth for measurement. The diameters of the culture were measured when mycelial growth on the corresponding untreated plates covered 75%-80% of the culture surface. The percentage of relative growth was calculated to determine the EC_50_ values. Two replicate plates were used for each treatment, and all experiments were performed two times.

### 2.3 Efficacy of ipflufenoquin on detached strawberry fruits and leaves

Five *B. cinerea* strains (CMML23-BC79, CMML23-BC74, CMML23-BC33, CMML23-BC14, and CMML23-BC95) were selected to test the efficacy of ipflufenoquin on strawberry fruits and leaves. These strains were selected based on mutations in the SDHI, QoI, and MBC fungicide target genes (*sdhB*, c*ytB*, and *tub2*, respectively). Fungicide sensitivity test results (EC_50_) are documented in the supplementary data (Table. S2). For the experiments, strawberry fruits and leaves were harvested from a greenhouse that has grown cultivar ‘Seolhyang’ and had not previously been exposed to fungicides. The surface of strawberry fruits and leaves was sterilized with 1% NaOCl for three minutes, rinsed three times with sterilized distilled water, and then air dried for 20 minutes. The commercial fungicide ipflufenoquin (Migiwa, Kyung Nong Co., Ltd., Korea, containing 20% of active ingredient) was sprayed using hand-held spray (applo) to the sterilized strawberry fruits and leaves at 12.5, 25, 50, 100, and 200 mg L^-1^ with 100 mg L⁻¹ being the recommended concentration. Three fungicide products – fluxapyroxad (Cardis, Nonghyup Chemical Co., Ltd., Korea, containing 15.3% of active ingredient), pyraclostrobin (Standby, Enbio Co., Ltd., Korea, containing 22.9% of active ingredient), and benomyl (FarmHannong Benomyl, FarmHannong, Co., Ltd., Korea, containing 50% of active ingredient) – were applied at 77, 67, and 250 mg L^-1^ concentrations, respectively. Water was sprayed as the control. The *B. cinerea* conidia were harvested from 7-day-old colonies, which were cultured under near-ultraviolet light at 12h/12h light-dark conditions at 25 °C. 1×10^5^ conidia mL^-1^ of conidia suspensions were prepared, and 10 μL of conidia suspensions was applied to each fruit, and a 5mm mycelial plug was inoculated onto each leaf. After incubating for three days at 25°C in the dark and 80% relative humidity, lesion sizes were measured with a digital caliper (Mitutoyo, Japan), and control efficacy was calculated relative to the fungicide-untreated control. Each experiment was performed four times.

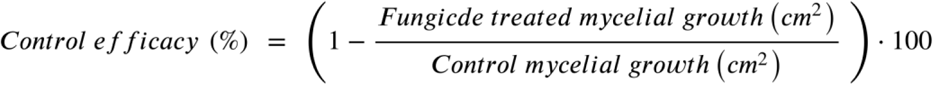

### 2.4 Control of gray mold in the greenhouse

A greenhouse experiment was conducted in February 2024 on strawberry plants in Gwangju, Korea. Plants which had been grown in a greenhouse for four months were treated with four spray applications. ipflufenoquin (Migiwa 20%) was applied at 25 and 100 mg L^-1^, and fluxapyroxad (Cardis 15.3%) was applied at 77 mg L^-1^ to the plants once a week for a total of three weeks. Plants in the control group were treated with water as the control. Following the only first application of the fungicide, a conidia suspension of *B. cinerea* strain CMML23-BC95, which showed sensitivity to MBC, QoI, and SDHI fungicides, was sprayed onto the plants for the ipflufenoquin efficacy test and CMML23-BC79 for the SDHI-resistant management test, respectively, at a concentration of 1×10^5^ conidia mL^-1^ and volume of 10 mL per plant. No further conidia suspension was applied in subsequent treatments. The control group was treated with water. To measure the results, the extent of disease occurrence on diseased fruits was investigated on days 14 and 21 after the fungicide treatment.

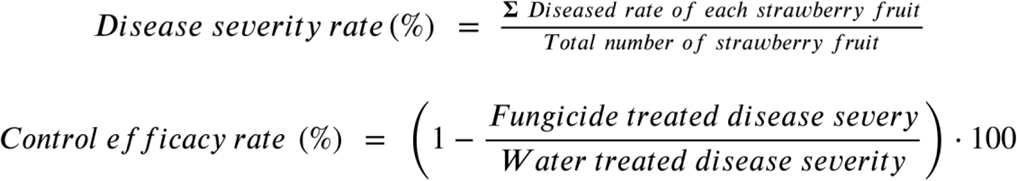

### 2.5 Postharvest fungal disease collection from the ipflufenoquin treatment plots

Four weeks after the first weekly application of ipflufenoquin, fungal diseases were collected from strawberry fruits. Fungal infections were investigated, and more than 30 fruits were harvested for each treatment of 25 and 100 mg L^-1^ of ipflufenoquin, and water only. Diseased tissues were detached and placed onto PDA agar plates, and a single colony was isolated from each plate after 3 days at 25℃ in the dark. Finally, one hundred ninety-seven fungal isolates were harvested from 25 mg L^-1^ (n = 93), 100 mg L^-1^ of ipflufenoquin (n = 74), and water only (n = 30).

### 2.6 In silico prediction of DHODH structure and molecular binding analysis

The structure of class 2 dihydroorotate dehydrogenase (DHODH), the target enzyme for ipflufenoquin, was predicted using Alphafold2 (Jumper et al., 2021). Amino acid sequences of DHODH were retrieved from the NCBI database: XP_018389332.1 for *Alternaria alternata*, QRD86425.1 for *A. flavus*, XP_024553432.1 for *B. cinerea*, XP_031876577.1 for *Colletotrichum fructicola*, XP_001223311.1 for *Chaetomium globosum*, XP_031045452.2 for *Fusarium oxysporum*, XP_011328027.1 for *Fusarium graminearum*, PHYSODRAFT_557375 for *P. sojae*, CEL63009.1 for *R. solani*, and XP_013952192.1 for *Trichoderma virens*. Additionally, DHODH nucleotide sequences were obtained from whole genome data of *Colletotrichum nymphaeae* strain JS-0361, *Colletotrichum truncatum* strain CMES1059, *Cladosporium tenuissimum* strain A3.I.1, and *Trichoderma gamsii* strain T6085. The sequence of *Rhizopus oryzae* Y5 strain was aligned more closely with Class 1A DHODH (Fig. S1), which uses fumarate as an electron acceptor instead of coenzyme Q10 (Oliver et al., 2016). Among the five predicted protein structures generated by Alphafold2, the structure with the highest confidence score was selected for subsequent binding studies. The 3D structure of ipflufenoquin and coenzyme Q10, used as the ligand, was obtained from the PubChem database (https://pubchem.ncbi.nlm.nih.gov/). Molecular docking was performed using AutoDock Vina v1.2.5 to estimate protein-ligand binding affinities (ΔG) (Eberhardt et al., 2021; Trott and Olson, 2010). For analysis, a grid box size of 30 was centered on the quinone binding site, which serves as the ligand’s target region. Each protein-ligand docking analysis was repeated 10 times for the 13 predicted protein structures. The highest binding affinities (-ΔG values) were statistically processed to calculate the mean and standard error. Finally, the protein-ligand binding affinities (ΔG) were compared to the LogEC_50_ values obtained from in vitro assays, and the results were visualized in a combined graph for comparative analysis.

## 3. Results and discussion

### 3.1 Diverse fungal species in strawberry plants and their potential pathogenic impact

The strawberry environment is surrounded by many fungal species, which may harbor potential pathogenicity for the plants and may originate from the root, stem, leaf, fruit, and rhizosphere (Li and Liu, 2019; Sangiorgio et al., 2022). The gray mold strains were collected from the fruits and leaves from 88 strawberry farms in Korea (Fig. 1), and the other 11 fungal genus groups were obtained from all parts of strawberry plants in the greenhouse (Table. S1). A total of 140 strains were identified as 7 strains of *A. alternata*, 88 of *B. cinerea*, 2 of *C. globosum*, 4 of *C. tenuissimum*, 4 of *C. fructicola*, 4 of *C. nymphaeae*, 3 of *Fusarium fujikuroi* species complex, 2 of *F. graminearum* species complex, 1 of *Fusarium incarnatum-equiseti* species complex, 20 of *F. oxysporum* species complex, 2 of *R. oryzae*, 1 of *Trichoderma atrobrunneum*, 1 of *T. gamsii*, 1 of *T. virens* (Table 1). Fungal isolates have been found to be involved in symptoms of fruit rot, black spot, anthracnose, crown rot, blossom blight, and wilt (Fu et al., 2020; Nam et al., 2015; Wedge et al., 2007; Zhang et al., 2022). In Korea, diverse fungal diseases have afflicted farms mainly with gray mold, crown rot, anthracnose, and powdery mildew (Je et al., 2015; Kim et al., 2023; Yoo et al., 2024). According to recent reports, strawberry crops are highly susceptible to several major fungal diseases, which significantly reduce yields and impact fruit quality. *B. cinerea* (gray mold), one of the most devastating pathogens, causes fruit rot that can lead to losses of up to 80%, particularly under cool, moist conditions. It affects primarily the flowers and fruits, causing water-soaked lesions that develop into gray moldy masses, rendering the fruit unmarketable (Nakielska et al., 2024). Similarly, blossom blight caused by *C. tenuissimum* affects strawberry production by infecting stigma of the strawberry flowers. It produces gray spores on the stigmas and causes necrosis across the entire flower, consequently inhibiting the normal formation of strawberry fruits. This disease occurs mainly within the Seolhyang cultivar and average disease incidence is 20% in greenhouses (Nam et al., 2015). Furthermore, Fusarium wilt, caused by *F. oxysporum* f. sp. *fragariae*, has posed a persistent threat to Korean strawberry crops since being first reported in the 1980s, leading to significant economic damages due to crop loss and also challenges associated with its management (Cho and Kwak, 2022). In this study, several fungal isolates were identified as major pathogens causing significant damage to strawberry cultivation and harvest in Korea. These findings highlight the critical need for integrated disease management strategies, including the development and use of resistant cultivars, effective fungicide applications, and improved cultural practices to mitigate the impact of these fungal pathogens. Korea’s high humidity in the rainy or winter season provides optimal growth conditions for fungi in the greenhouse. Gray mold disease is one of the most destructive and rapidly spreading infections in fruits and flowers when there is high humidity with cool temperatures, so this fungus was targeted mainly for disease control in this study.

**Figure 1.**
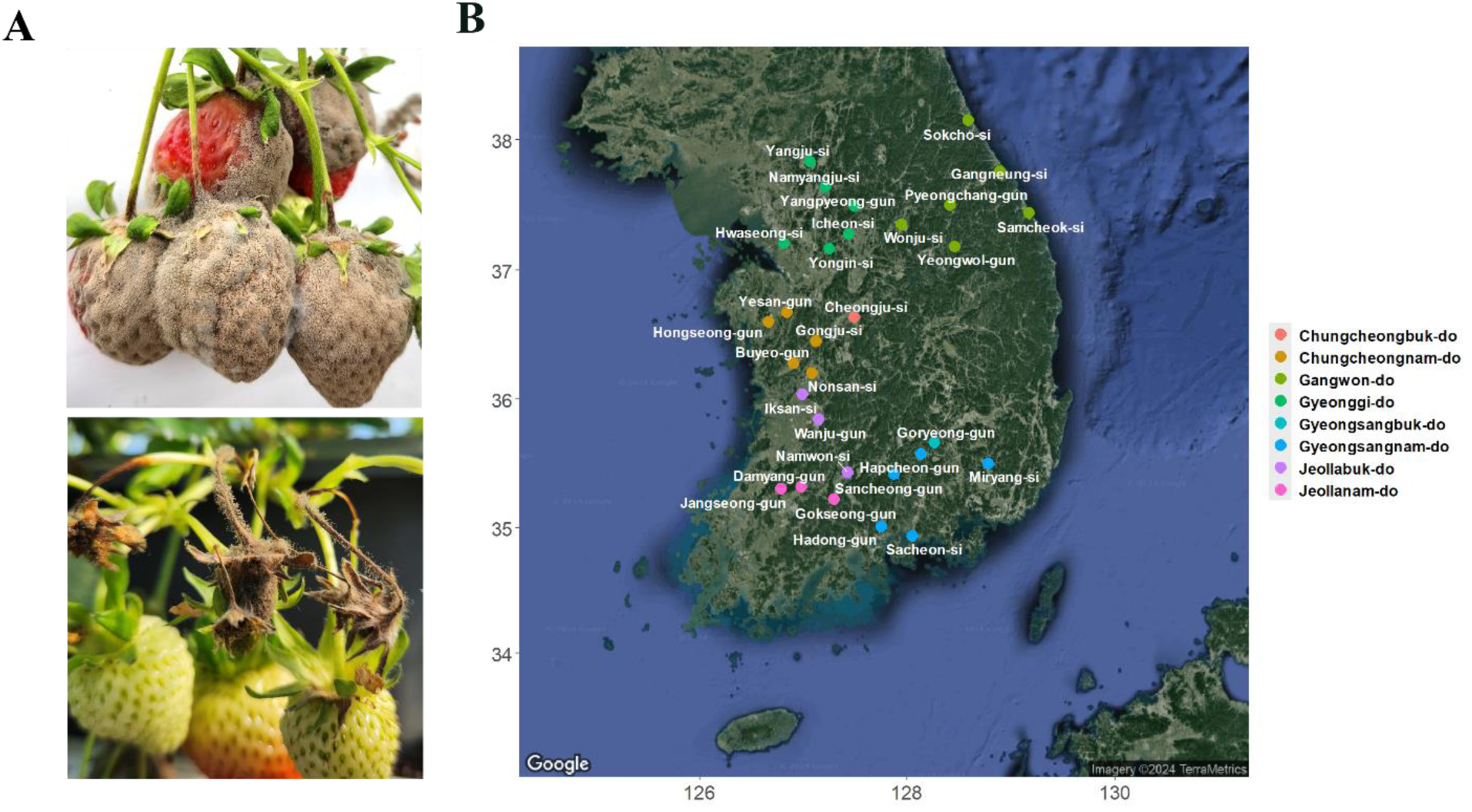
Isolation of *Botrytis cinerea*. Gray mold disease was observed on strawberry fruits and flowers during the spring season from March to May in Korea (A). A map showed locations from which *B. cinerea* 88 isolates were obtained (B). Each color dot represents strains(n=1∼3) that were collected in 2023.

**Table 1.** Application of fungicides against various phytopathogenic fungi. Fungicide susceptibility tests were performed on pathogenic fungi collected from various environments, including strawberries. EC_50_ values greater than 10 are considered less susceptible and are highlighted in red.

| Fungal species<br>(Group) | Number<br>of isolates | Host<br>(source) | Primers for<br>identification | EC<sub>50</sub> values<br>(mg·L<sup>-1</sup>) | Media |
| --- | --- | --- | --- | --- | --- |
| *Botrytis cinerea* | 88 | Strawberry | *ITS, NEP2* | 0.0025-2.88 | PDA |
| *Alternaria alternata* | 7 | Strawberry | *ITS, ALT1* | 0.04-1.05 | PDA |
| *Chaetomium globosum* | 2 | Strawberry | *ITS, EF1* | 0.0016-0.0018 | PDA |
| *Cladosporium tenuissimum* | 4 | Strawberry | *ITS, DHODH* | >100 | PDA |
| *Colletotrichum fruticola* | 4 | Strawberry | *ITS, GAPDH* | 0.005-0.077 | PDA |
| *Colletotrichum nymphaeae* | 4 | Strawberry | *ITS, GAPDH* | 0.24-0.31 | PDA |
| *Fusarium* spp. (FIESC) | 1 | Strawberry | *ITS, TUB2* | 0.69 | PDA |
| *Fusarium* spp. (FGSC) | 2 | Strawberry | *ITS, TUB2* | 0.08-0.14 | YBA |
| *Fusarium* spp. (FFSC) | 3 | Strawberry | *ITS, TUB2* | 0.01-0.04 | PDA |
| *Fusarium* spp. (FOSC) | 20 | Strawberry | *ITS, TUB2* | 0.09-0.88 | PDA |
| *Rhizopus oryzae* | 2 | Strawberry | *ITS* | 30.98-40.95 | PDA |
| *Trichoderma atrobrunneum* | 1 | Strawberry | *ITS, EF1* | 0.27 | PDA |
| *Trichoderma gamsii* | 1 | Strawberry | *ITS, EF1* | 0.33 | PDA |
| *Trichoderma virens* | 1 | Strawberry | *ITS, EF1* | 0.7 | PDA |
| *Aspergillus flavus* | 2 | Meju, soil | *ITS, CYP51* | 0.002-0.005 | PDA |
| *Colletotrichum truncatum* | 2 | Soybean | *ITS* | 0.08-0.11 | PDA |
| *Phytophthora sojae* | 1 | Soybean | *ITS* | 17.28 | CMA |
| *Rhizoctonia solani* AG1 | 3 | Rice | *ITS* | >100 | PDA |
| *R. solani* AG-2-2 (III B) | 1 | bentgrass | *ITS* | >100 | PDA |

### 3.2 EC*_50_* distribution of Botrytis cinerea and various phytopathogenic fungal strains to ipflufenoquin

The ipflufenoquin susceptibility test on *B. cinerea* strains showed that the EC_50_ value for mycelial growth lay between 0.0025 and 2.88 mg L^-1^ (Fig. 2A) Fungicide action was more effective on conidial elongation, ranging from 0.0095 to 0.41 mg L^-1^ (Fig. 2B). The average EC_50_ values were 0.46±0.07 mg L^-1^ and 0.07±0.027 mg L^-1^ for each. Although there was no significant correlation in EC_50_ values between mycelial growth and conidia elongation, fungicidal activity on growth rate was effective in both. No resistance was found in any *B.cinerea* strain. This study provides the first baseline sensitivity test for *B. cinerea* isolates collected from strawberries across various regions using ipflufenoquin, offering important insights into the current sensitivity levels of the pathogen and serving as a reference point for future resistance monitoring and management efforts.

**Figure 2.**
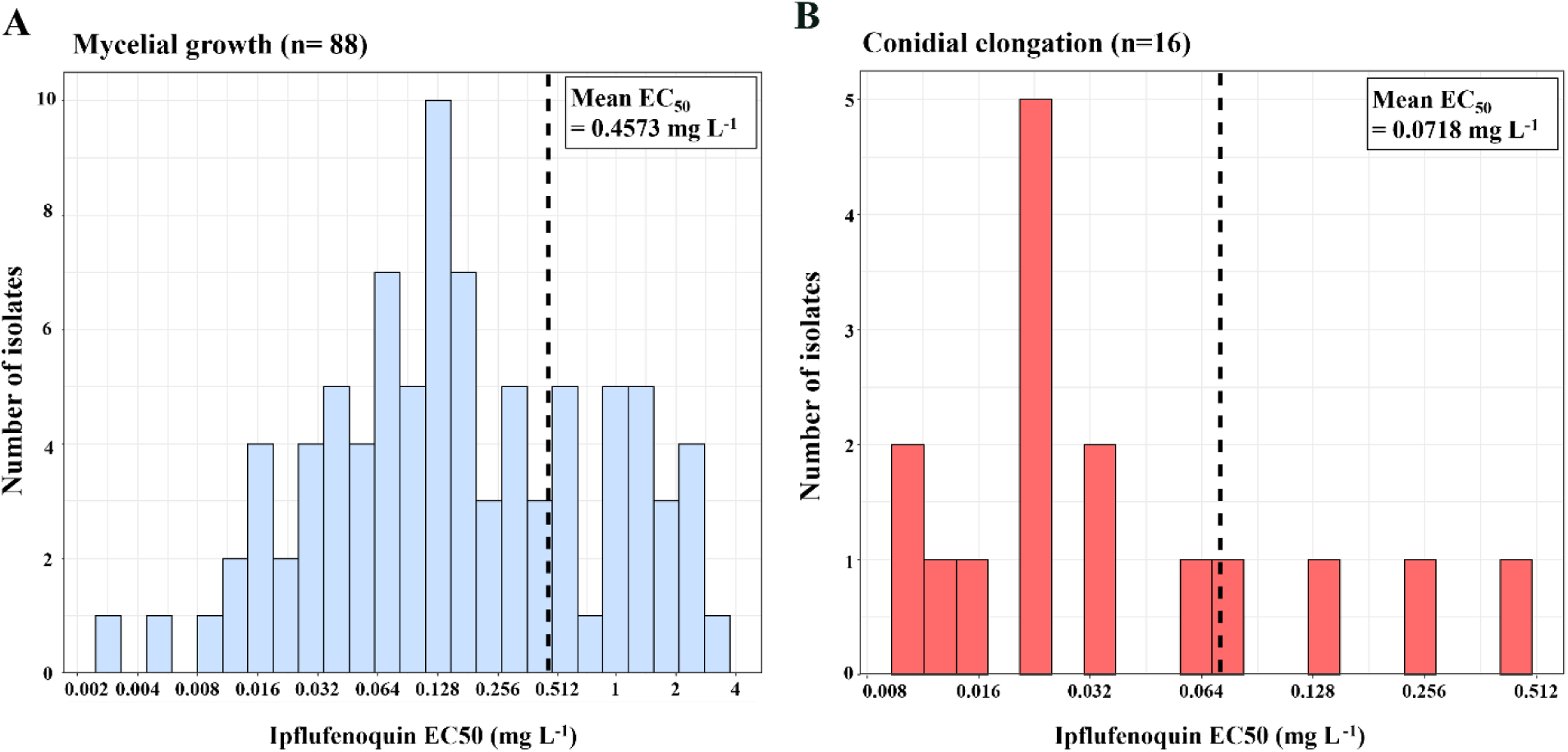
Frequency distribution of fungicide concentrations resulting in 50% growth inhibition (EC_50_) to ipflufenoquin. EC_50_ values were measured for each sample, using mycelium (A) and conidia (B), along with the mean EC_50_. The LL.3 model of the R package “drc” was used to calculate EC_50_.

Among the fungi isolated from strawberries, *C. tenuissimum* and *R. oryzae* showed EC_50_ values exceeding 100 mg L^-1^, indicating highly reduced sensitivity to ipflufenoquin; *A. alternata* showed EC_50_ values ranging from 0.04 mg L^-1^ to over 100 mg L^-1^. The other fungal species showed susceptible phenotypes to ipflufenoquin (0.0007–1.69 mg L^-1^) (Table. 1). In the case of fungi from other hosts, *A. flavus* (0.002-0.005 mg L^-1^, n = 2), *C. truncatum* (0.08–0.11 mg L^-1^, n = 2), and *F. graminearum* species complex (0.06-0.2 mg L^-1^, n = 8) showed susceptible phenotypes. But *R. solani* (>100 mg L^-1^, n = 4) and *P. sojae* (17.28 mg L^-1^, n = 1) proved resistant to ipflufenoquin. From the perspective of taxonomic classification, Basidiomycota, Mucoromycota, and Oomycete were insensitive to ipflufenoquin, as well as some of the Ascomycota species; *A. alternata* (3 of 13 isolates), and *C. tenuissium* showed insensitivity as high EC_50_ values (> 10 mg L^-1^) (Table. 1). The DHODH inhibitor, ipflufenoquin, can be applied to most mold diseases, but it could be challenging to control all fungal species in plant pathogens. As a similar DHODH inhibitor, olorofim has shown significant efficacy in sensitivity tests against various fungal species, particularly *A. fumigatus*, including azole-resistant strains, with consistently low MIC values (Buil et al., 2017). However, its effectiveness against other fungi varied in one study with higher MICs observed for species like *F. solani* and dematiaceous fungi such as *A. alternata* and *Exophiala dermatitidis*, indicating reduced potency against these groups (Feuss et al., 2024). Additionally, recent research has shown that strains of *A. fumigatus* exposed to the agricultural fungicide ipflufenoquin can develop cross-resistance to olorofim. The study demonstrated that mutations in the DHODH gene lead to high resistance levels against both compounds, with ipflufenoquin exhibiting a higher IC_50_ compared to olorofim, indicating that it is less effective in inhibiting the enzyme than olorofim (Van Rhijn et al., 2023). The current study confirmed the specific sensitivity to ipflufenoquin by various plant pathogenic fungi, including those affecting strawberries, providing relevant information that can be utilized to enhance disease control measures and resistance monitoring in agricultural practices.

### 3.3 Effect of gray mold control on strawberry fruits and leaves by fungicides

The effect of fungicide on strawberry fruits was observed with four products: ipflufenoquin, fluxapyroxad, pyraclostrobin, and benomyl (Fig. 3A). The growth inhibition range of ipflufenoquin was 87%–98%, demonstrating its effectiveness against all strains of *B. cinerea* (Fig. 3B). On the other hand, fruits treated with fluxapyroxad, pyraclostrobin, and benomyl, were susceptible to multiple fungicide resistant strains with the SDHB-P225F, CYTB-G143A, and TUB2-G198V mutations (CMML23-BC33, CMML23-BC74, and CMML23-BC79). Also, the CMML23-BC14 strain, which has CYTB-G143A and TUB2-G198V mutations, showed insensitivity to pyraclostrobin and benomyl. The CMML23-BC95 strain, which does not contain any fungicide target genes mutations, was highly controlled by fluxapyroxad and benomyl (90%, and 83% efficacy), and potency was greatly reduced in pyraclostrobin (39% efficacy). All the fungicide-resistant strains in this study, which the chemicals SDHI, QoI, and MBC could not control, were effectively managed by ipflufenoquin in the strawberry fruits. Ipflufenoquin can be a good alternative antifungal agent when conventional fungicides must be applied many times to strawberry plants. To check the efficacy of low concentrations of a new DHODH inhibitor fungicide in the fruit, ipflufenoquin was sprayed at six different concentrations (12.5 to 200 mg L^-1^). The results showed that the mycelial growth of five *B. cinerea* strains was significantly inhibited by all concentrations (83% to 98% efficacy) compared to the untreated control (Fig. 3C). The leaf assay using mycelial plugs displayed diminished efficacy, with 49% to 77% in 100 mg L^-1^ ipflufenoquin (Fig. S2). The mycelial plug exhibited a higher pathogenic infection on leaves. Pre-treatment of ipflufenoquin before conidia germination would represent good timing to manage gray mold disease.

**Figure 3.**
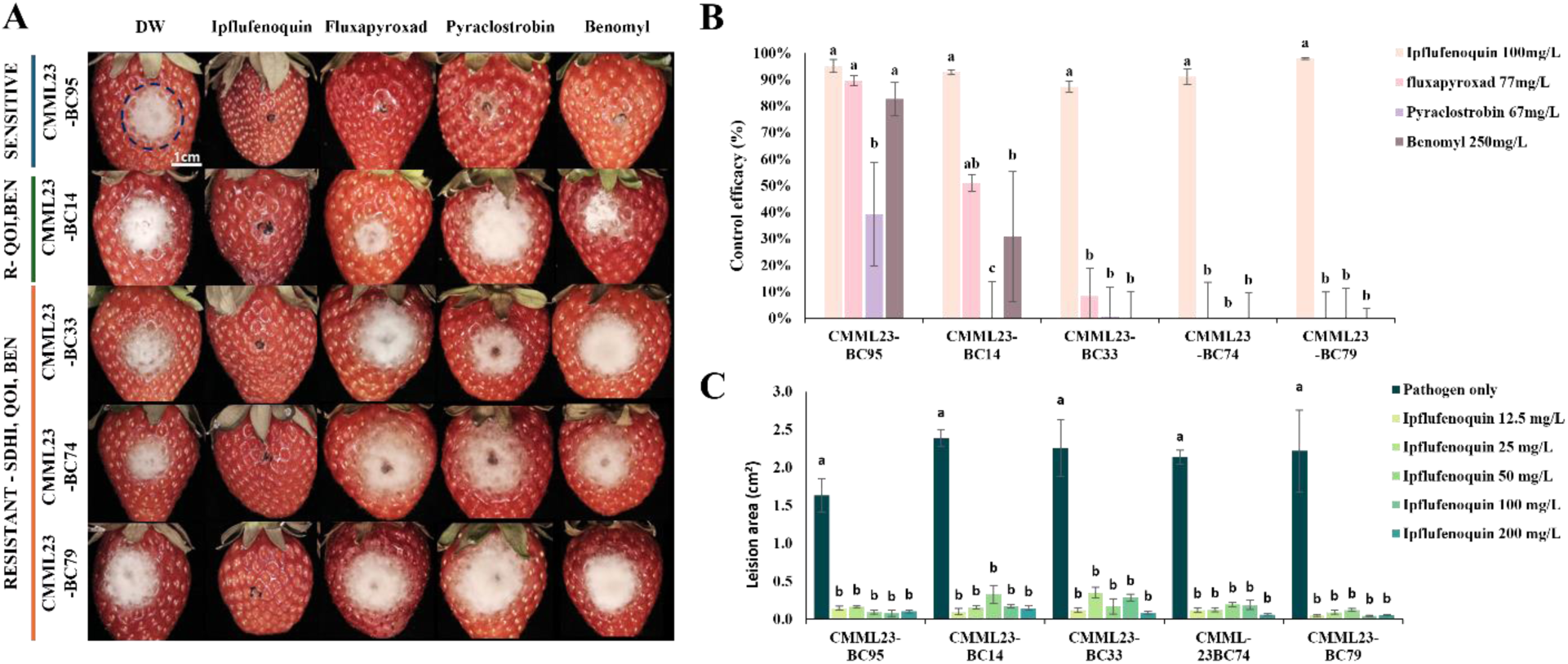
Fungicide efficacy tests of *B. cinerea* on strawberry fruits. Strains with mutations in *sdhB*, *cytB*, or *tub2* genes (CMML23-BC14, CMML23-BC33, CMML23-BC74, CMML23-BC79) and a non-mutated strain (CMML23-BC95) were selected and inoculated on fruits treated with various fungicides (A). Control efficacy for each strain was calculated by comparison with fungicide-untreated strawberries (B). A range of ipflufenoquin concentrations (12.5 to 200 mg L⁻¹) was applied to evaluate its detailed control effectiveness (C).

Managing multiple fungicide-resistant populations is crucial for maintaining effective disease control and preventing significant agricultural losses. Resistance to multiple fungicides, as seen in pathogens like *B. cinerea* and *Fusarium* spp., reduces treatment options and diminishes the efficacy of control measures, leading to higher costs and environmental impacts (De Mio et al., 2024) However, the application of ipflufenoquin offers a promising solution, as it can help reduce fungicide-resistant populations. Even at low concentrations, such as 12.5 mg L⁻¹, ipflufenoquin is expected to provide sufficient disease control in strawberries, further promoting its role in integrated disease management strategies.

### 3.4 Fungicide efficacy tests in greenhouse

Application of ipflufenoquin (25 and 100 mg L^-1^) to strawberry plants in a greenhouse was performed once a week, up to 3 times. No symptoms were observed until 7 days, but from 14 days post-inoculation (dpi), the CMML23-BC95 strain showed typical symptoms of gray mold, with 17±5.7% disease severity, which was increased up to 79±6.0% at 21 dpi (Fig. 4A). The effectiveness of 100 mg L^-1^ ipflufenoquin (recommended concentration by the company) was 100% until 14 days and was gradually reduced to 94% after an additional week. On the other hand, the efficacy of 25 mg L^-1^ was decreased by 81% on the 14^th^ day and 65% on the 21^st^ day. (Fig. 4B). When treated with 100 mg L^-1^ of ipflufenoquin, flowers showed a distinct control effect compared to the untreated group. (Fig. 4D). When greenhouse conditions are conducive for gray mold infection with low temperatures (2∼10℃) and high humidity, a weekly application of 100 mg L^-1^ of ipflufenoquin could be recommended for long-term disinfection.

**Figure 4.**
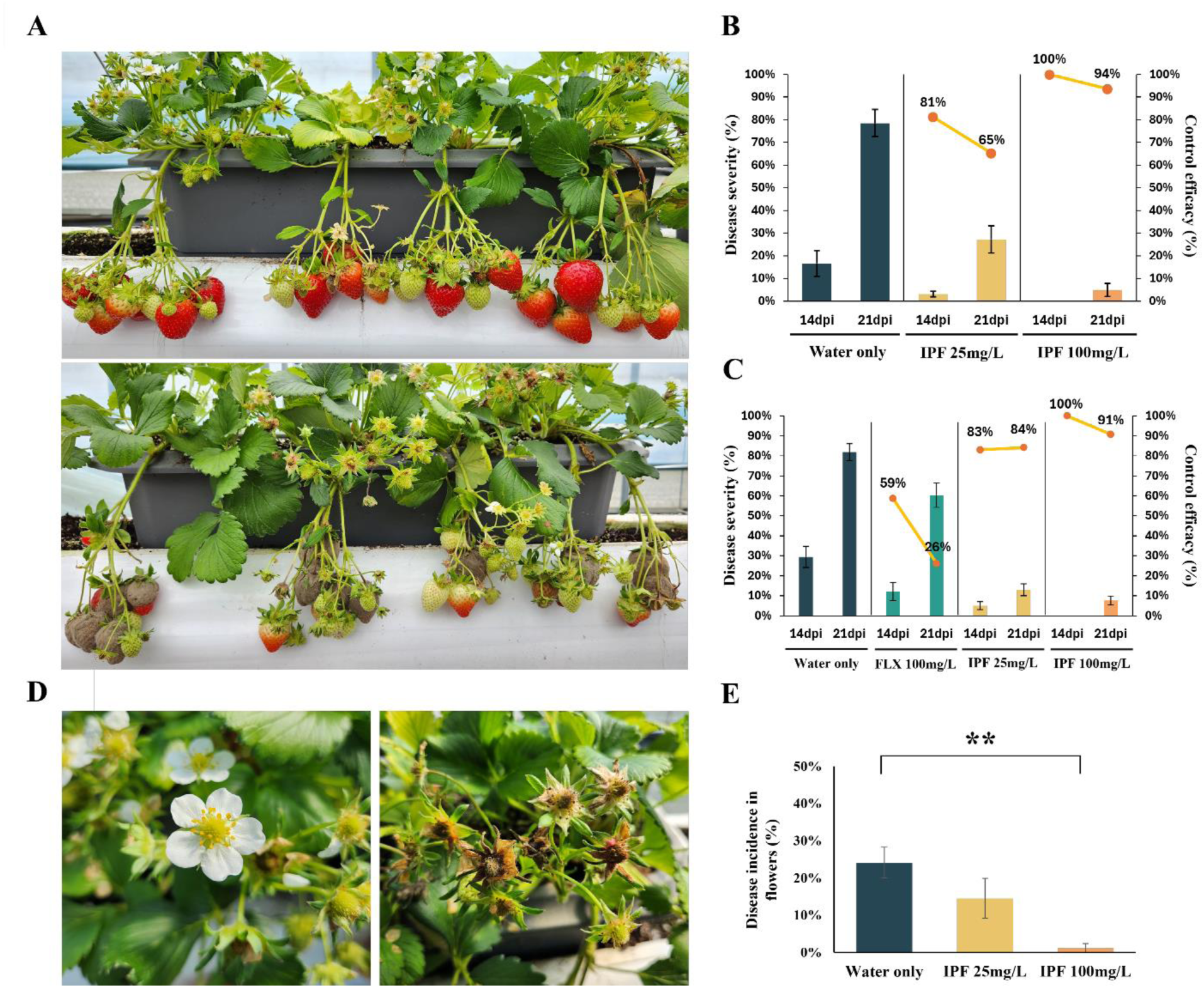
Control of gray mold disease by ipflufenoquin treatment in a greenhouse. Ipflufenoquin at 100 mg/L was applied to plants inoculated with *B. cinerea* strain CMML23B-C95 (A, top), while only the pathogen was applied in the control group (A, bottom). Disease severity and control efficacy were assessed for the sensitive strain CMML23-BC95 (B) and the fluxapyroxad-resistant strain CMML23-BC79 (C). The efficacy of ipflufenoquin was evaluated on flowers 3 weeks post-treatment, comparing treated (D, left) and untreated (D, right) flowers, with efficacy compared in (E). *IPF* = ipflufenoquin, *FLX* = fluxapyroxad.

The SDHI-resistant strain CMML23-BC79 (SDHB-P225F) was tested against fluxapyroxad and the ipflufenoquin. Efficacy of fluxapyroxad was 59% in 14 dpi and 28% in 21 dpi. Gray mold with the P225F mutation in *sdhB* was not sufficiently controlled when fluxapyroxad was used. On the other hand, ipflufenoquin controlled the fungal strain, showing 83% to 100% of control efficacy when treated with 25 and 100 mg L^-1^ (Fig. 4C). Recent studies have reported that more than 70% of *B. cinerea* isolates collected in Korea during 2020-2021 exhibited resistance to the SDHI fungicide boscalid, with identified mutations in the *sdhB* gene, such as P225F, P225H, and H272R. (Liu et al., 2024). Based on field reports from strawberry crops in northern Germany, *B. cinerea* isolates have exhibited high levels of resistance to several fungicides, including the quinone-outside inhibitors (fenhexamid, boscalid, and cyprodinil) with resistance rates surpassing 20% (Weber and Hahn, 2019). Additionally, the study of Diánez et al. (2002) from Huelva, Spain reported a 94% benomyl resistance in *B. cinerea* isolates from strawberry fields, while the study of Kim et al. (2023) in Korea found that 82.05% of isolates were resistant, primarily due to E198A and E198V mutations in the β-tubulin gene. These results suggest that benomyl resistance has persisted over two decades, driven by continuous benzimidazole fungicide use, highlighting the need for carefully selected fungicide selection and rotation to manage resistance and ensure effective disease control across various regions (Diánez et al., 2002; Kim et al., 2023). The current study is the first to explore the use of ipflufenoquin for controlling both natural and SDHI-resistant strains of *B. cinerea* in strawberry fields. It provides practical recommendations for field application and suggests that ipflufenoquin could effectively manage fungicide-resistant populations.

### 3.5 The occurrence of various other fungal diseases

Upon completion of the control experiment in March 2024, naturally occurring gray mold was dominant in the fungicide-untreated area. Gray mold disease was effectively suppressed with the application of ipflufenoquin. Nonetheless, various pathogenic molds such as blossom blight (*Cladosporium* spp.), Rhizopus rot (*Rhizopus* spp.), and black spot (*Alternaria* spp.) were relatively not suppressed (Fig. 5A). Increasing the ipflufenoquin treatment concentration decreased the amount of gray mold disease. Still, the proportion from 39.8% to 47.3% of blossom blight and from 14% to 18.9% of Rhizopus rot was increased (Fig. 5B). Later tests on strawberries showed that *Cladosporium* spp. from the ipflufenoquin 100 mg L⁻¹ treatment group did not respond to the chemical (Fig. S3). These results indicate that in untreated plots without selection pressure, gray mold can rapidly spread across strawberry fields without other fungal pathogens. However, as the concentration of ipflufenoquin increased, an undesirable side effect was observed, with populations of *Rhizopus* (from Mucoromycota) and *Cladosporium* (from Ascomycota) appearing more frequently. The fungal composition in strawberries, including persistent species like *B. cinerea*, may be altered by using fungicides such as ipflufenoquin. This mirrors findings from a study showing that fungicide treatments could lead to the dominance of more resistant species, such as *Basidiomycota*, in pesticide-treated soils (Streletskii et al., 2022). This indicates that secondary infections or new plant diseases could emerge as resistant species capitalize on the vacated ecological niches of more sensitive fungi.

**Figure 5.**
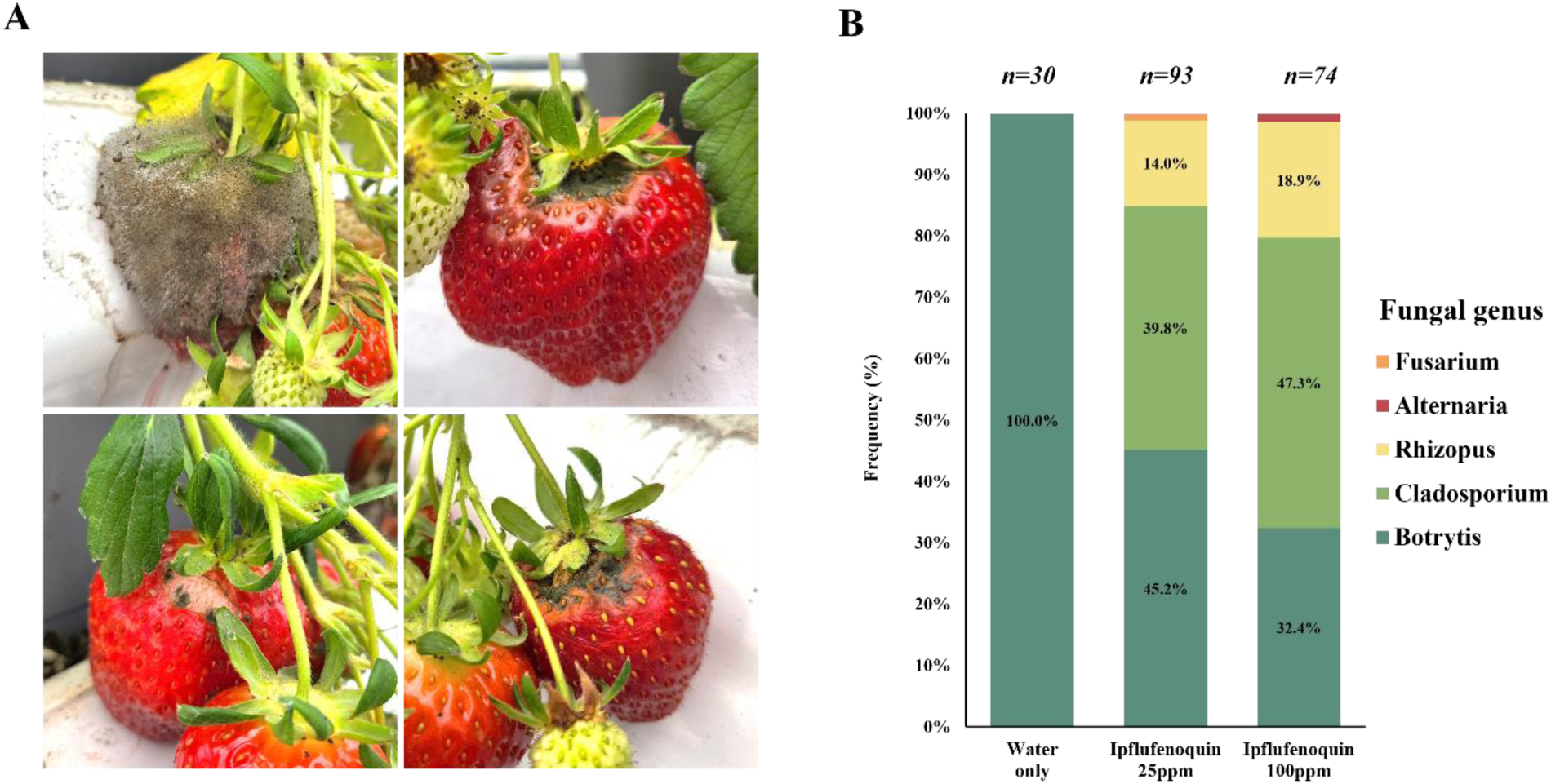
Occurrence of various other fungal diseases. After 4 weeks of ipflufenoquin treatment (long-term use), the occurrence of various fungal diseases was observed, and mold sampling was conducted (A). According to morphological characteristics, various genus of fungi such as *Cladosporium, Rhizopus, Alternaria,* and *Fusarium*, as well as *Botrytis*, were isolated (B).

### 3.6 Dihydroorotate dehydrogenase (DHODH) structure prediction and binding study of Ipflufenoquin

The protein structure of *B. cinerea* DHODH was predicted using AlphaFold2, revealing that both coenzyme Q10 and ipflufenoquin are located within the quinone binding tunnel. The calculated binding affinities were -8.518 kcal mol⁻¹ for coenzyme Q10 and -9.604 ± 0.008 kcal mol⁻¹ for ipflufenoquin (Fig. 6A,B). The alpha-helices (ɑA, ɑB, and ɑD) and beta-sheets (βC and βD) were identified within 5Å of the ligand, ipflufenoquin, following the protein structure classification by Hansen et al., 2004 (Fig. 6C). Among the DHODH protein structures of the 13 fungal species, *C. globosum* exhibited the highest binding affinity at -9.875 ± 0.004 kcal mol⁻¹, whereas *R. solani* showed the lowest at -7.523 ± 0.076 kcal mol⁻¹ (Fig. 7B). A graph was generated to compare binding affinities (ΔG) with in vitro fungicide susceptibility data (logEC_50_ values). *C. globosum* and *A. flavus*, which had EC_50_ values below 0.003 mg L⁻¹, exhibited protein-ligand binding affinities of less than -9.684 kcal mol⁻¹, while *C. tenuissimum* and *R. solani*, which had EC_50_ values above 100 mg L⁻¹, showed binding affinities greater than -8.221 kcal mol⁻¹ (Fig. 7C).

**Figure 6.**
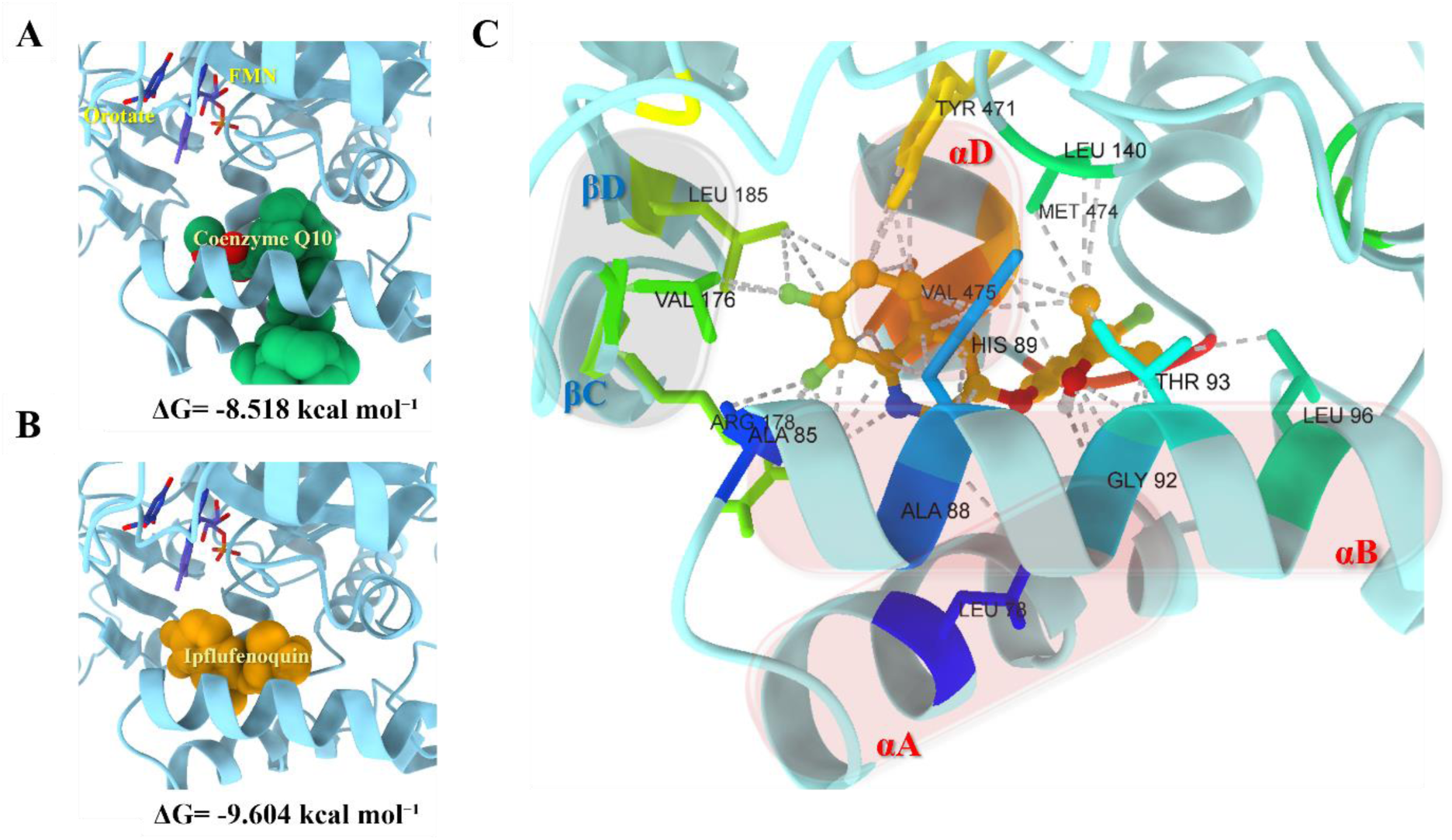
Structure prediction of *B. cinerea* dihydroorotate dehydrogenase (DHODH). The binding simulation with Coenzyme Q10 (A) and ipflufenoquin (B). Three alpha-helices and two beta-sheets are located adjacent to ipflufenoquin in the quinone-binding tunnel (C).

**Figure 7.**
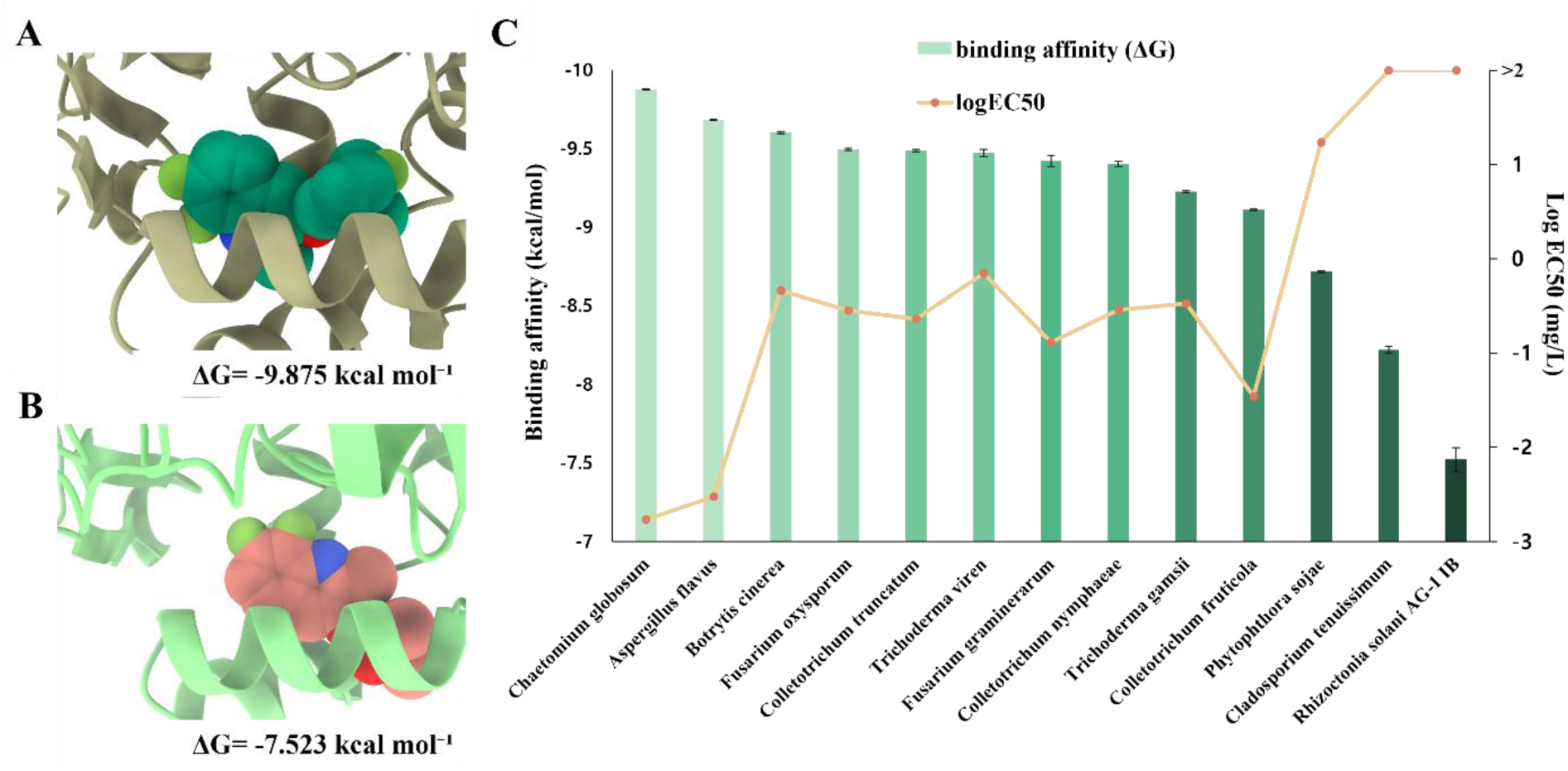
Binding simulation and affinity evaluation of fungal DHODH to ipflufenoquin. The lowest binding affinity (ΔG) was observed in *Chaetomium globosum* DHODH (A); the highest, in *Rhizoctonia solani* DHODH (B). Binding affinities (ΔG) and fungicide sensitivity values (LogEC_50_) are presented for 13 fungal species included in the study (C).

Interaction of ipflufenoquin with the quinone binding tunnel disrupts the electron acceptor function of coenzyme Q10, thereby inhibiting subsequent metabolic processes (Boschi et al., 2019). Based on the crystal structures of other known DHODH inhibitors, such as atovaquone, brequinar, and teriflunomide, it has been hypothesized that their binding sites are similar to that of ipflufenoquin (Hansen et al., 2004; Liu et al., 2000). These inhibitors interact with important structural components near the quinone binding tunnel, specifically the three alpha-helices and two beta-sheet, which are crucial for effective inhibition (Fig. S4). Studies on DHODH inhibitors have demonstrated that mutations in these structural regions significantly affect fungicide susceptibility, as recently observed in Aspergillus fumigatus (Van Rhijn et al., 2023). The DHODH structures in the 13 fungal species aligned with the general mechanism of Class 2 DHODH, and their functions are expected to follow this mechanism as well. However, different amino acid sequences were identified near the quinone binding tunnel across species, and these variations are thought to influence the binding affinity of DHODH inhibitors (Fig. S5). This study suggests a general correlation between binding affinity and in vitro fungicide susceptibility results, providing a useful foundation for developing models to better predict fungicide susceptibility across fungal species.

## 4. Conclusion

This study offers significant insights into the efficacy of ipflufenoquin against *B. cinerea* and other fungal pathogens associated with strawberry cultivation. Field experiments demonstrated that ipflufenoquin exhibited strong control over both natural and SDHI-resistant strains of *B. cinerea*, even at low concentrations, underscoring its potential as an effective alternative to conventional fungicides. Its novel mode of action, targeting DHODH, suggests that its application may help reduce the emergence of resistance, improving long-term disease control in the agricultural environment. The action of ipflufenoquin may lead to unintended changes in the composition of pathogenic fungal communities in strawberries, as the reduction of *B. cinerea* may create an opportunity for fungi less sensitive to fungicides, such as *C. tenuissimum* and *R. oryzae* to proliferate, potentially resulting in new plant health issues. Binding affinity analysis further clarifies the relationship between ipflufenoquin’s interaction with fungal DHODH and the corresponding in vitro fungicide susceptibility. Fungal species exhibiting higher binding affinities, such as *C. globosum* and *A. flavus*, demonstrated increased sensitivity to ipflufenoquin. In contrast, species like *C. tenuissimum* and *R. solani*, which had lower binding affinities, displayed reduced sensitivity. These results establish a basis for developing predictive models that can accurately assess fungicide susceptibility across a wide range of fungal pathogens, ultimately supporting more precise and effective disease management strategies.

## CRediT authorship contribution statement

Sungyu Choi: Conceptualization, Writing - Original Draft, Methodology, Software, Visualization. Doeun Son: Writing – Review & Editing, Methodology, Formal Analysis, Data Curation, Visualization. Hyeong-rok Jang: Writing – Review & Editing, Investigation, Data Curation. Soyoon Park: Writing – Review & Editing, Investigation, Data Curation. Haifeng Liu: Methodology, Software, Investigation, Data Curation. Hyeon-gyeong Lee: Investigation. Hyunkyu Sang: Conceptualization, Writing – Review & Editing, Project Administration, Funding Acquisition.

## Data Availability

Data will be made available on request.

## Acknowledgments

This work was supported by the Rural Development Administration (Republic of Korea), grant number PJ01690602.

## Supplementary data

**Table S1.** *Botrytis cinerea* strains used in this study.

| No | Strain name | Location | GPS Coordinates |
| --- | --- | --- | --- |
| 1 | CMML23-BC01 | Sacheon-si, Gyeongsangnam-do | 35°6'25.99055455846667" N, 127°54'57.335587160512205" E |
| 2 | CMML23-BC02 | Sacheon-si, Gyeongsangnam-do | 35°6'27.53780053667981" N, 127°54'56.671879698351404" E |
| 3 | CMML23-BC03 | Sacheon-si, Gyeongsangnam-do | 35°7'49.15847317657153" N, 127°55'17.408699499776503" E |
| 4 | CMML23-BC04 | Hadong-gun, Gyeongsangnam-do | 35°12'46.24884598270455" N, 127°52'50.46498740910465" E |
| 5 | CMML23-BC05 | Hadong-gun, Gyeongsangnam-do | 35°11'31.32578857957867" N, 127°52'46.95215076317709" E |
| 6 | CMML23-BC06 | Hadong-gun, Gyeongsangnam-do | 35°12'44.87227992688531" N, 127°53'11.682955743181083" E |
| 7 | CMML23-BC08 | Hapcheon-gun, Gyeongsangnam-do | 35°33'14.895662021642124" N, 128°14'37.76368998639782" E |
| 8 | CMML23-BC10 | Hapcheon-gun, Gyeongsangnam-do | 35°33'38.4300276880208" N, 128°15'16.01863285170566" E |
| 9 | CMML23-BC11 | Miryang-si, Gyeongsangnam-do | 35°22'5.661219665036583" N, 128°43'33.59051196299106" E |
| 10 | CMML23-BC12 | Miryang-si, Gyeongsangnam-do | 35°22'11.37052784177854" N, 128°43'30.55584058099157" E |
| 11 | CMML23-BC13 | Miryang-si, Gyeongsangnam-do | 35°22'16.790669915336593" N, 128°43'38.23002617544489" E |
| 12 | CMML23-BC14 | Miryang-si, Gyeongsangnam-do | 35°22'23.788859360799393" N, 128°43'28.51940620283358" E |
| 13 | CMML23-BC15 | Goryeong-gun, Gyeongsangbuk-do | 35°50'47.20431732923771" N, 128°24'50.07566428610971" E |
| 14 | CMML23-BC16 | Goryeong-gun, Gyeongsangbuk-do | 35°50'45.15180439660469" N, 128°24'49.93075689767238" E |
| 15 | CMML23-BC17 | Goryeong-gun, Gyeongsangbuk-do | 35°50'44.80729684237019" N, 128°24'45.56138166021583" E |
| 16 | CMML23-BC18 | Cheongdo-gun, Gyeongsangbuk-do | 35°34'26.244254671960334" N, 128°47'31.535045433399773" E |
| 17 | CMML23-BC19 | Cheongdo-gun, Gyeongsangbuk-do | 35°34'33.2104909825523" N, 128°47'38.40358924322118" E |
| 18 | CMML23-BC20 | Cheongdo-gun, Gyeongsangbuk-do | 35°34'38.12147179892861" N, 128°47'43.017093052678774" E |
| 19 | CMML23-BC21 | Namwon-si, Jeollabuk-do | 35°21'51.29032191691692" N, 127°20'56.28510064286161" E |
| 20 | CMML23-BC22 | Namwon-si, Jeollabuk-do | 35°21'42.98584843427818" N, 127°20'33.418565405162326" E |
| 21 | CMML23-BC24 | Gokseong-gun, Jeollanam-do | 35°17'29.011976770987076" N, 127°18'57.618612060970236" E |
| 22 | CMML23-BC25 | Gokseong-gun, Jeollanam-do | 35°17'12.719297243266965" N, 127°18'53.759557777187865" E |
| 23 | CMML23-BC26 | Gokseong-gun, Jeollanam-do | 35°18'3.090020943672016" N, 127°18'57.345420634881066" E |
| 24 | CMML23-BC27 | Damyang-gun, Jeollanam-do | 35°20'55.05609750061069" N, 126°59'12.318190166636214" E |
| 25 | CMML23-BC28 | Damyang-gun, Jeollanam-do | 35°21'23.047229416914092" N, 126°59'41.12767044499378" E |
| 26 | CMML23-BC29 | Damyang-gun, Jeollanam-do | 35°15'19.875856648826584" N, 126°55'28.546973963653954" E |
| 27 | CMML23-BC30 | Jangseong-gun, Jeollanam-do | 35°14'45.672683197248034" N, 126°50'41.8420044370464" E |
| 28 | CMML23-BC31 | Jangseong-gun, Jeollanam-do | 35°14'58.05118297651916" N, 126°50'42.53427144937518" E |
| 29 | CMML23-BC33 | Iksan-si, Jeollabuk-do | 36°4'3.1720532074791663" N, 127°4'18.762949308487578" E |
| 30 | CMML23-BC34 | Iksan-si, Jeollabuk-do | 36°4'14.20324827932177" N, 127°3'57.175445500037085" E |
| 31 | CMML23-BC35 | Iksan-si, Jeollabuk-do | 36°2'58.47416619794956" N, 127°1'31.881861687301125" E |
| 32 | CMML23-BC36 | Wanju-gun, Jeollabuk-do | 35°56'46.9422390529013" N, 127°9'42.267485483345126" E |
| 33 | CMML23-BC37 | Wanju-gun, Jeollabuk-do | 35°54'42.677828564817446" N, 127°6'15.666060715769845" E |
| 34 | CMML23-BC38 | Wanju-gun, Jeollabuk-do | 35°54'39.62907037654929" N, 127°5'56.1296845252798" E |
| 35 | CMML23-BC39 | Cheongju-si, Chungcheongbuk-do | 36°32'24.389275965059483" N, 127°30'46.83785127497799" E |
| 36 | CMML23-BC41 | Cheongju-si, Chungcheongbuk-do | 36°43'19.94472885233847" N, 127°32'43.62796463496352" E |
| 37 | CMML23-BC42 | Yesan-gun, Chungcheongnam-do | 36°41'30.352331266049646" N, 126°46'5.190797012110124" E |
| 38 | CMML23-BC43 | Yesan-gun, Chungcheongnam-do | 36°39'56.82743097171738" N, 126°38'32.66298081835657" E |
| 39 | CMML23-BC44 | Yesan-gun, Chungcheongnam-do | 36°41'17.084562442277615" N, 126°49'8.817956058456957" E |
| 40 | CMML23-BC45 | Buyeo-gun, Chungcheongnam-do | 36°18'9.156935973701934" N, 126°51'58.005764577929995" E |
| 41 | CMML23-BC46 | Buyeo-gun, Chungcheongnam-do | 36°11'49.933952656407996" N, 126°44'56.507733135700846" E |
| 42 | CMML23-BC50 | Gongju-si, Chungcheongnam-do | 36°23'0.8008662577890391" N, 127°4'19.434093160967905" E |
| 43 | CMML23-BC51 | Gongju-si, Chungcheongnam-do | 36°19'44.24544479928784" N, 127°7'37.12089981901158" E |
| 44 | CMML23-BC52 | Nonsan-si, Chungcheongnam-do | 36°10'55.876036168569954" N, 127°7'55.72342932442666" E |
| 45 | CMML23-BC53 | Nonsan-si, Chungcheongnam-do | 36°7'14.20309203429781" N, 127°6'8.376420744181132" E |
| 46 | CMML23-BC54 | Nonsan-si, Chungcheongnam-do | 36°10'28.998220314582568" N, 127°7'54.68624132959576" E |
| 47 | CMML23-BC55 | Nonsan-si, Chungcheongnam-do | 36°10'27.793427082772837" N, 127°7'54.68624132959576" E |
| 48 | CMML23-BC56 | Hongseong-gun, Chungcheongnam-do | 36°33'28.944051160426625" N, 126°34'12.589620803075832" E |
| 49 | CMML23-BC57 | Hongseong-gun, Chungcheongnam-do | 36°39'48.260508653499414" N, 126°42'44.57755034342654" E |
| 50 | CMML23-BC58 | Hongseong-gun, Chungcheongnam-do | 36°40'4.613811472438556" N, 126°43'52.23919700900524" E |
| 51 | CMML23-BC59 | Samcheok-si, Gangwon-do | 37°24'3.186119736472506" N, 129°12'32.366295207582425" E |
| 52 | CMML23-BC60 | Samcheok-si, Gangwon-do | 37°23'37.33392841908028" N, 129°12'58.822909489849735" E |
| 53 | CMML23-BC61 | Samcheok-si, Gangwon-do | 37°24'11.430818138916834" N, 129°12'37.210696842980724" E |
| 54 | CMML23-BC62 | Sokcho-si, Gangwon-do | 38°11'39.665197939026484" N, 128°34'5.09385341145844" E |
| 55 | CMML23-BC63 | Sokcho-si, Gangwon-do | 38°10'54.16598856453163" N, 128°33'6.061932456664181" E |
| 56 | CMML23-BC64 | Sokcho-si, Gangwon-do | 38°10'57.53006057920231" N, 128°33'13.188028647443844" E |
| 57 | CMML23-BC65 | Gangneung-si, Gangwon-do | 37°50'59.47519093378446" N, 128°45'50.09439812586152" E |
| 58 | CMML23-BC66 | Gangneung-si, Gangwon-do | 37°51'17.61627757719225" N, 128°50'28.107028600970807" E |
| 59 | CMML23-BC67 | Gangneung-si, Gangwon-do | 37°50'59.125584735636494" N, 128°51'26.92860574437873" E |
| 60 | CMML23-BC68 | Yeongwol-gun, Gangwon-do | 37°11'41.72291841375625" N, 128°33'54.56694089089751" E |
| 61 | CMML23-BC69 | Yeongwol-gun, Gangwon-do | 37°10'40.140621991028524" N, 128°26'28.906990413249787" E |
| 62 | CMML23-BC70 | Yeongwol-gun, Gangwon-do | 37°17'39.08113918996776" N, 128°14'49.68336566621247" E |
| 63 | CMML23-BC71 | Wonju-si, Gangwon-do | 37°13'26.89487012372183" N, 128°5'3.802140894231343" E |
| 64 | CMML23-BC72 | Wonju-si, Gangwon-do | 37°13'7.960707227784951" N, 128°5'22.871791121474416" E |
| 65 | CMML23-BC73 | Wonju-si, Gangwon-do | 37°13'10.560521369903313" N, 128°5'20.95299803791022" E |
| 66 | CMML23-BC74 | Pyeongchang-gun, Gangwon-do | 37°26'42.285402858314" N, 128°25'55.48135711545683" E |
| 67 | CMML23-BC75 | Pyeongchang-gun, Gangwon-do | 37°26'39.311840176187616" N, 128°25'42.79572568779031" E |
| 68 | CMML23-BC76 | Pyeongchang-gun, Gangwon-do | 37°36'46.81855041885058" N, 128°22'2.0559171395734666" E |
| 69 | CMML23-BC77 | Chuncheon-si, Gangwon-do | 37°55'48.436174511650165" N, 127°46'32.72662289957907" E |
| 70 | CMML23-BC78 | Chuncheon-si, Gangwon-do | 37°54'20.17720239088362" N, 127°43'51.414540989527495" E |
| 71 | CMML23-BC79 | Chuncheon-si, Gangwon-do | 37°55'48.352603006964046" N, 127°46'31.529387515186045" E |
| 72 | CMML23-BC80 | Icheon-si, Gyeonggi-do | 37°10'42.74469461317949" N, 127°30'40.118243744376514" E |
| 73 | CMML23-BC81 | Icheon-si, Gyeonggi-do | 37°10'59.4481305431799" N, 127°30'49.437453270098786" E |
| 74 | CMML23-BC82 | Icheon-si, Gyeonggi-do | 37°13'22.788516549492215" N, 127°28'53.69097994121262" E |
| 75 | CMML23-BC83 | Yangpyeong-gun, Gyeonggi-do | 37°32'8.72720434547773" N, 127°18'50.5709895101927" E |
| 76 | CMML23-BC84 | Yangpyeong-gun, Gyeonggi-do | 37°32'49.80513666565571" N, 127°19'35.796758083652094" E |
| 77 | CMML23-BC85 | Yangpyeong-gun, Gyeonggi-do | 37°32'14.902051329365236" N, 127°18'57.564709509939576" E |
| 78 | CMML23-BC86 | Hwaseong-si, Gyeonggi-do | 37°13'25.342652456042174" N, 127°2'17.42181327610865" E |
| 79 | CMML23-BC87 | Hwaseong-si, Gyeonggi-do | 37°10'6.259654598403586" N, 127°4'43.92916469640568" E |
| 80 | CMML23-BC88 | Hwaseong-si, Gyeonggi-do | 37°14'3.172659883293818" N, 127°3'5.745845659096176" E |
| 81 | CMML23-BC89 | Yongin-si, Gyeonggi-do | 37°10'58.824359812189186" N, 127°18'4.302660888891978" E |
| 82 | CMML23-BC90 | Yongin-si, Gyeonggi-do | 37°19'47.07759122379173" N, 127°14'28.34050948172603" E |
| 83 | CMML23-BC91 | Yongin-si, Gyeonggi-do | 37°8'58.68079716552302" N, 127°12'7.68058945648022" E |
| 84 | CMML23-BC92 | Yangju-si, Gyeonggi-do | 37°47'34.82477045591736" N, 127°6'2.2003647827739314" E |
| 85 | CMML23-BC93 | Yangju-si, Gyeonggi-do | 37°47'24.6382314987261" N, 127°5'27.672972401398965" E |
| 86 | CMML23-BC95 | Namyangju-si, Gyeonggi-do | 37°33'52.908838226268244" N, 127°19'5.802125704830132" E |
| 87 | CMML23-BC96 | Namyangju-si, Gyeonggi-do | 37°33'48.6837779050785" N, 127°19'6.694614276606217" E |
| 88 | CMML23-BC97 | Namyangju-si, Gyeonggi-do | 37°34'4.7904829703958285" N, 127°19'12.94951046779488" E |

**Table S2.**
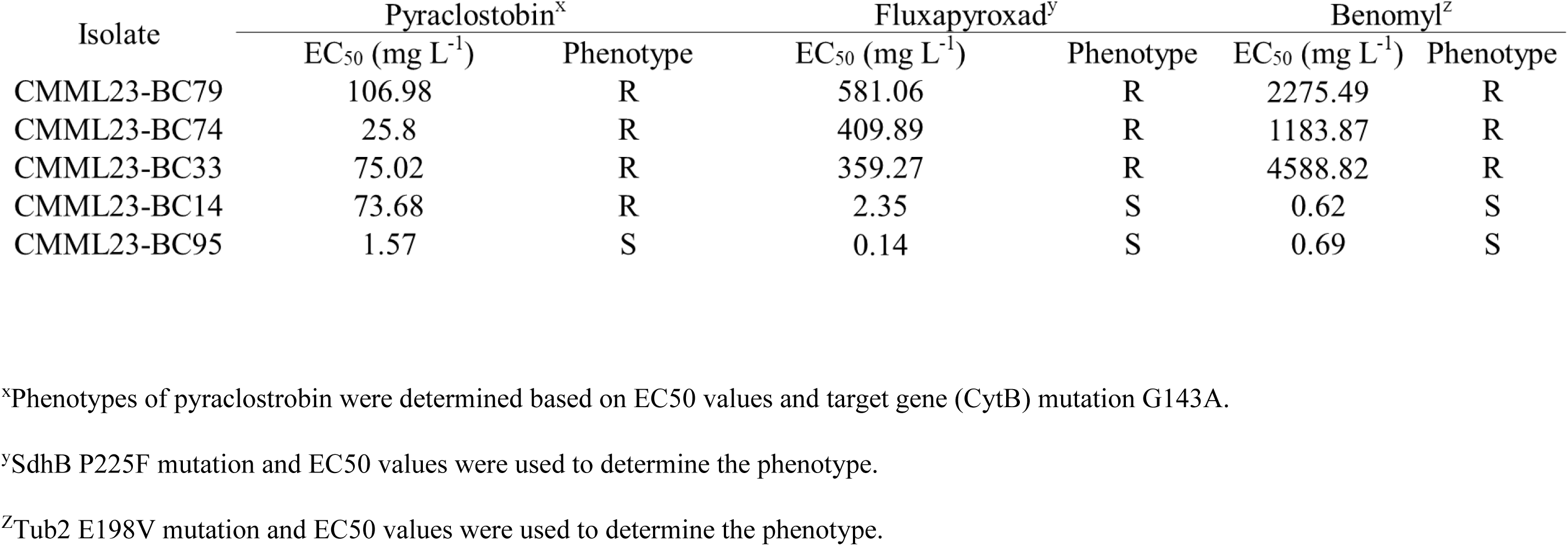
Fungicide sensitivity test to *B. cinerea* strains in this study.

**Figure S1.**
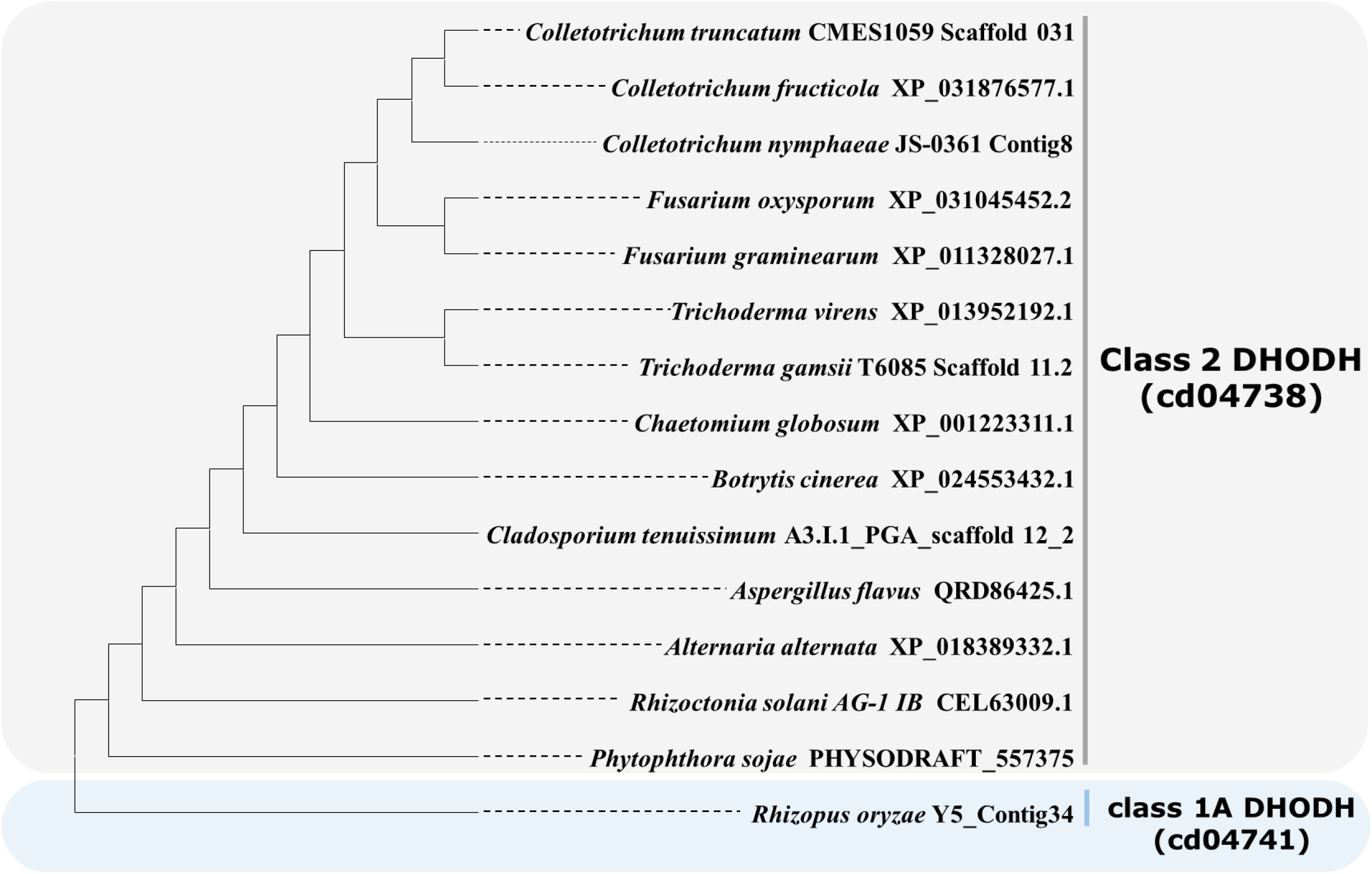
Phylogenetic analysis of dihydroorotate dehydrogenase (DHODH). The DHODH sequences were aligned, and the tree was generated using maximum likelihood method in Mega X program.

**Figure S2.**
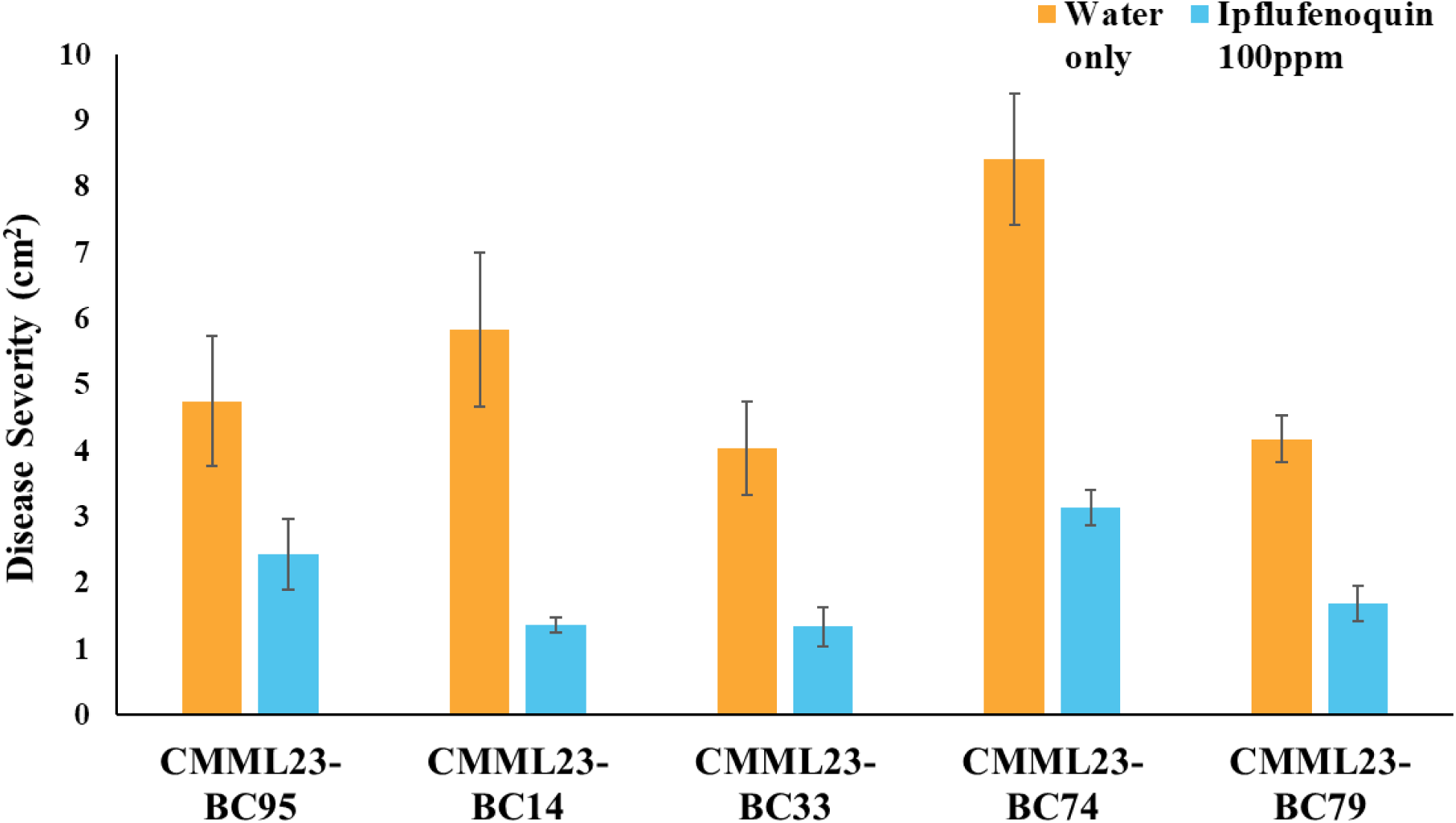
Fungicide efficacy tests of *B. cinerea* on strawberry leaves. Mycelial growth, the disease severity, was reduced when ipflufenoquin was applied to the leaves.

**Figure S3.**
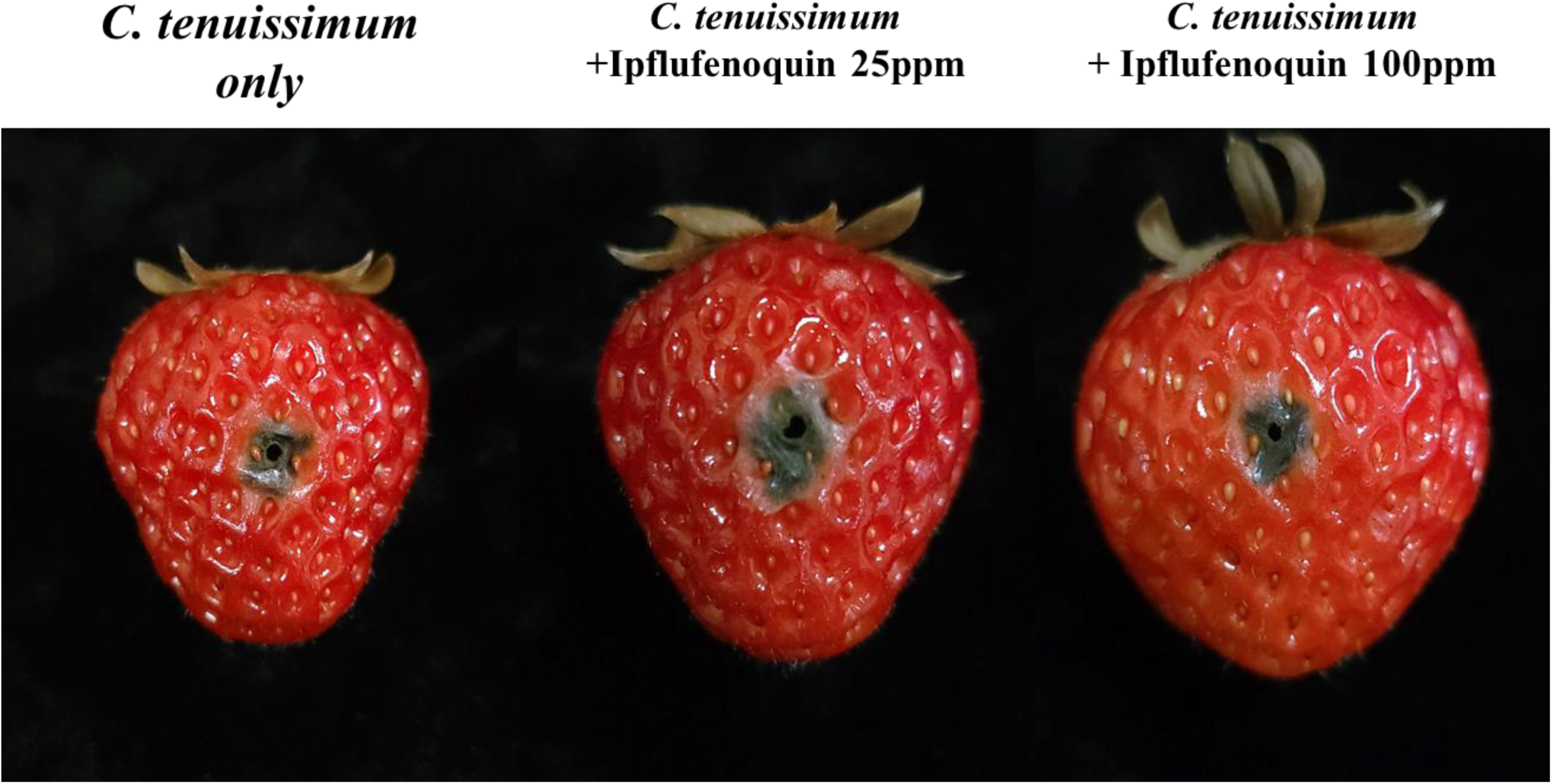
Fungicide susceptibility of *Cladosporium tenuissimum* isolated from strawberries. Strawberries were inoculated with spores to test the fungicide effect. It appears to have minimal control effect compared to other pathogenic fungi.

**Figure S4.**
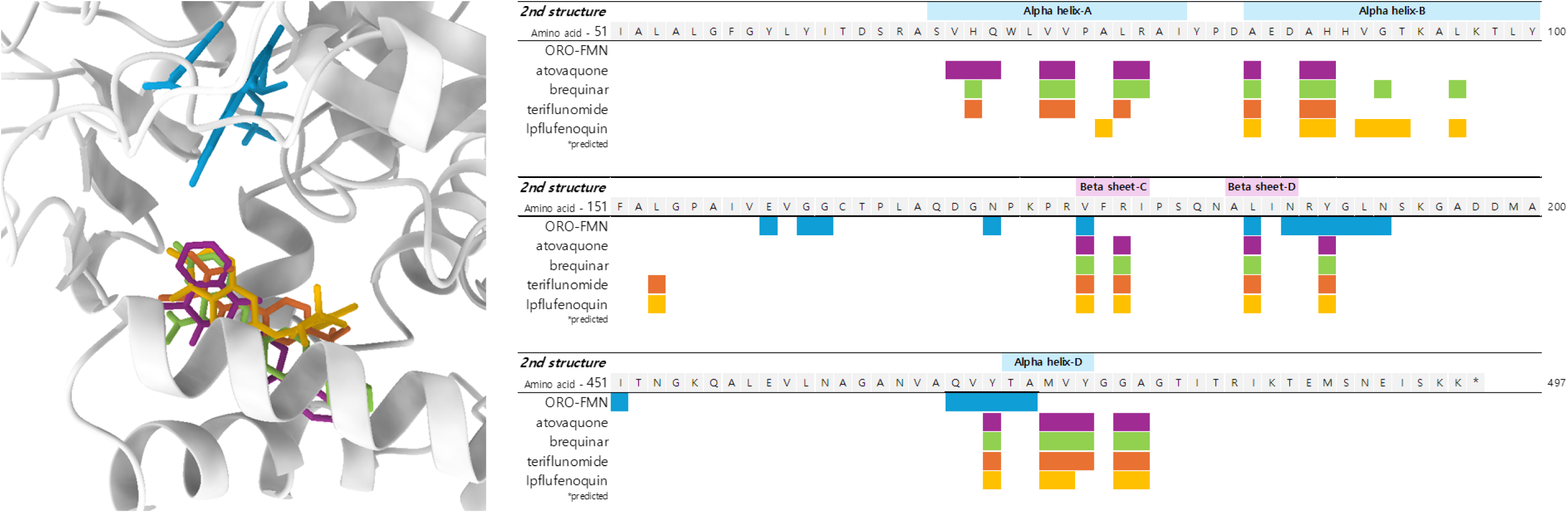
Binding sites of DHODH inhibitors in *B. cinerea* DHODH and adjacent amino acids. *B. cinerea* DHODH structure is shown in gray, with orotate (ORO) and FMN in blue, atovaquone in purple, brequinar in green, teriflunomide in orange, and ipflufenoquin (predicted) in dark yellow.

**Figure S5.**
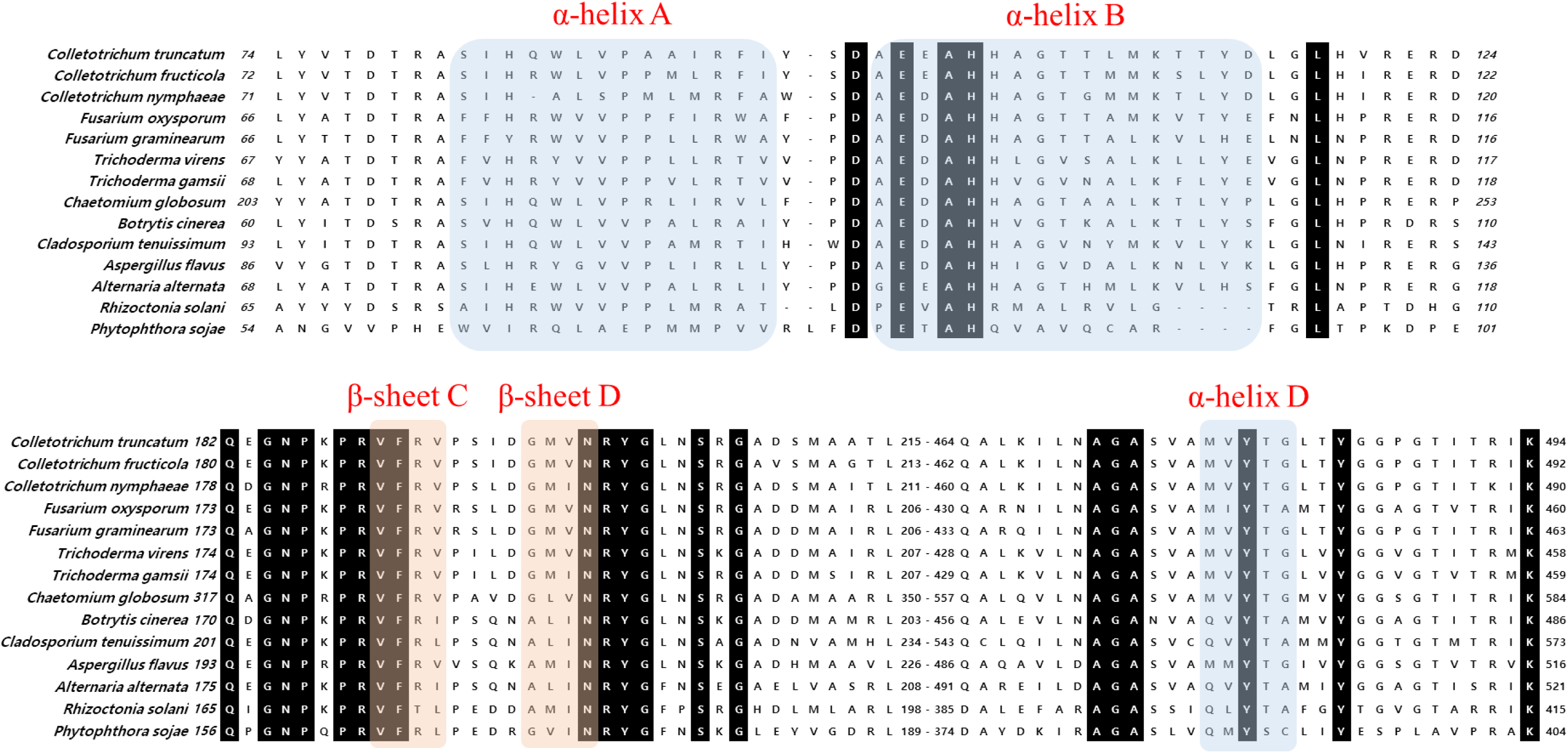
Amino acid sequence comparison of three alpha-helices and two beta-sheets within the quinone-binding tunnel of DHODH across 13 fungal species. Alpha-helices are shown in blue; beta-sheets in apricot.

